# Accelerating long-read analysis on modern CPUs

**DOI:** 10.1101/2021.07.21.453294

**Authors:** Saurabh Kalikar, Chirag Jain, Vasimuddin Md, Sanchit Misra

## Abstract

Long read sequencing is now routinely used at scale for genomics and transcriptomics applications. Mapping of long reads or a draft genome assembly to a reference sequence is often one of the most time consuming steps in these applications. Here, we present techniques to accelerate minimap2, a widely used software for mapping. We present multiple optimizations using SIMD parallelization, efficient cache utilization and a learned index data structure to accelerate its three main computational modules, i.e., seeding, chaining and pairwise sequence alignment. These result in reduction of end-to-end mapping time of minimap2 by up to 1.8 × while maintaining identical output.

## Background

Long-read or single molecule sequencing technology from Pacific Biosciences (PacBio) and Oxford Nanopore Technology (ONT) have made significant leaps in terms of read lengths, sequencing throughput and accuracy since their introduction to the market. Longer read lengths naturally benefit genomics and transcriptomics applications, e.g., to detect complex structural variation in case of DNA sequencing [1], or for novel isoform discovery during RNA sequencing [2]. As a result, long read sequencing is now being adopted in population-scale and biodiversity genome surveys [3–5]. However, increased sequencing throughput, e.g., *>* 1 Tbp per day [6], also demands faster processing of data to save time and cloud computing costs. Among the many steps performed to analyse a long-read data set, mapping of long DNA or RNA reads to a reference sequence is usually the first and among the most time-consuming steps in any bioinformatics workflow.

Minimap2 is a widely used sequence alignment program which supports many use-cases including mapping long reads or a draft genome assembly to a reference sequence [7]. Even though minimap2 uses well-engineered heuristics and software libraries, its performance remains significantly below peak computing performance of a modern CPU. Presence of frequent branching in the code, irregular memory accesses and irregular computation in minimap2 make it challenging to efficiently utilise the available hardware resources. Owing to its complexity, only a few attempts have been made to accelerate minimap2, and they have also been confined to accelerating only one of the three modules within minimap2 [8–10]. The highest speed-up reported till date for minimap2 on multicore CPUs remains 1.4x, and this was achieved without guaranteeing identical output as the original implementation [10].

### Minimap2 Algorithm

The algorithm used in minimap2 [7] is based on the standard seed-chain-align procedure (Figure 1). The seeding stage identifies short fixed-length exact matches between a read and a reference sequence. Minimap2 makes use of minimizer technique [11], a popular k-mer sampling method to improve time and space requirements. Prior to mapping, minimap2 does offline indexing of the reference sequence where it builds a multimap using a hash table with minimizers as keys and minimizer locations as values. This hash table is used during the seeding step when exact matches are collected by searching read minimizers in the reference index. Such matching pairs of minimizers form a set of anchors which are sorted and passed onto the chaining stage. From the complete list of sorted anchors, the chaining stage identifies an ordered subset of anchors that are co-linearly positioned along a diagonal [12, 13]. Minimap2 uses a customized chaining score function to prioritise the highest-scoring chains which are likely to yield the desired base-to-base alignments of a read. It uses dynamic programming for chaining and has two versions: DP chaining and RMQ-based DP chaining. DP chaining algorithm is *O*(*n*^2^) in the number of anchors and is used when the number of anchors are expected to be small. RMQ-based DP chaining algorithm is *O*(*nlog*(*n*)) in the number of anchors and is used when the number of anchors are expected to be large. It is used as a “long-join” heuristic in minimap2 to chain anchors that are too far in the array. It uses a simplified cost function whereas DP chaining penalizes gaps more effectively. The third and final alignment stage computes base-level alignments for filling the gaps between adjacent anchors in these chains.

**Figure 1.**
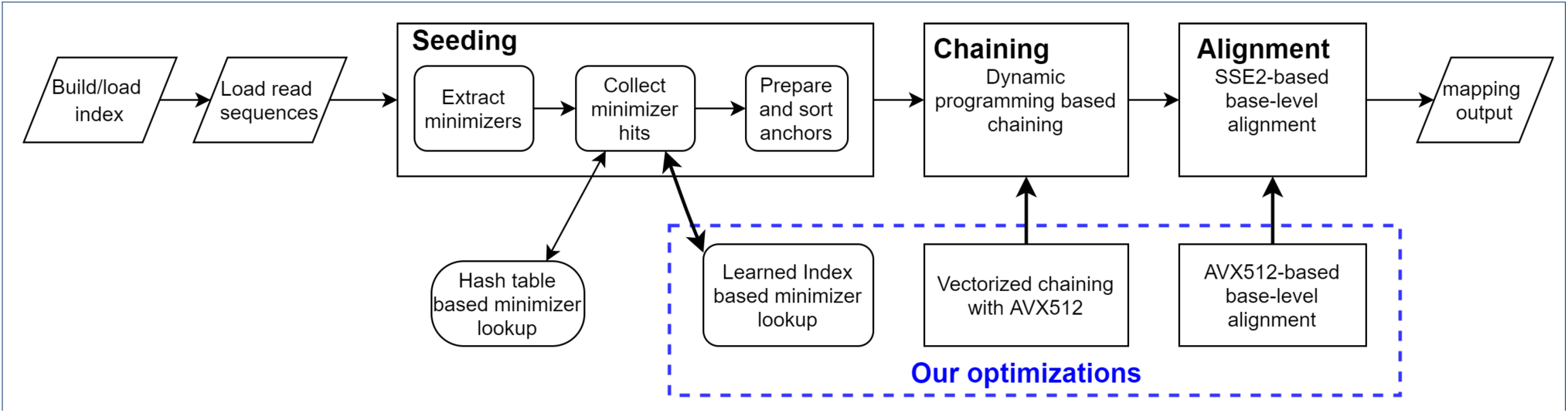
Minimap2 workflow depicting its three key modules: (i) seeding, (ii) chaining, and (iii) alignment. Our optimizations to each of the modules are shown in the blue dotted rectangle.

### Our contributions

In this work, we re-engineered the three key computational modules in minimap2: (i) seeding, (ii) anchor chaining, and (iii) pairwise sequence alignment. Optimization of the seeding stage was achieved by replacing the standard hash-table lookup with a machine learning based lookup using an hardware-efficient implementation of learned index data structure [14]. Acceleration of the anchor chaining step was achieved by designing a novel SIMD-parallel chaining algorithm which uses vector processing units available on modern CPUs. In the final sequence alignment stage, we reduced runtime by converting 128-bit (SSE) SIMD instructions to 256-bit (AVX2) and 512-bit (AVX512) SIMD instructions. In all the proposed optimizations, we ensured that the final output remains 100% identical to that of minimap2, which allows users to easily switch to a faster version of minimap2 whenever faster computing throughput is desired.

We compared our optimised minimap2 implementation *mm2-fast* with minimap2 by mapping real ONT, PacBio CLR, PacBio HiFi human sequencing data, and human de novo genome assemblies to the human genome reference using multiple generations of server grade CPUs. For each of the three modules, we developed both AVX2 and AVX512 based versions. Given the wider SIMD width, our AVX512 version achieves significantly higher speedups compared to the AVX2 version. The speedups achieved by mm2-fast AVX512 version ranged from 1.7 − 1.8×, 1.4 − 1.7×, 1.5 − 1.6×, and 1.3 − 1.5× for ONT, PacBio CLR, PacBio HiFi and genome-assembly inputs respectively.

To the best of our knowledge, no prior work has reported better speedup of end-to-end runtime of minimap2 using a single processor or even using hsingle CPU, single GPUi or hsingle CPU, single FPGAi combinations. mm2-fast implementation is open-sourced and available at https://github.com/lh3/minimap2/tree/fast-contrib-v2.22.

## Results

In this section, we discuss our experimental settings, various datasets we used, a summary of optimizations implemented in mm2-fast, and finally demonstrate the performance gains by comparing mm2-fast against minimap2.

### Experimental setup

We performed our experiments on four different processor architectures, Intel^®^ Xeon^®^ Platinum 8180 (Skylake), Intel^®^ Xeon^®^ Platinum 8280 (Cascade Lake), Intel^®^ Xeon^®^ Platinum 8380 (Ice Lake), and AMD EPYC™ 7742 (Rome). Architectural specifications of these system are listed in Supplementary Table S1. Our implementation mm2-fast was built on top of minimap2 (v2.22), therefore, all our benchmarks show a comparison of mm2-fast with minimap2 (v2.22). Our tests involved three types of real human long-read sequencing data (ONT Guppy 3.6.0, PacBio HiFi, PacBio CLR), and also three human genome assemblies for mapping to the standard reference GRCh38 [15]. The latter was useful to demonstrate utility of mm2-fast for faster genome-to-genome comparisons. Long-read sequencing datasets used here were available publicly, and derived from human trio benchmark genomes HG002, HG003 and HG004 (Table 1). The three human genome assemblies are associated with nearly-haploid CHM13 [16] and diploid HG002 genomes [17]. Each type of dataset was mapped using parameters recommended in minimap2 documentation.

**Table 1.** Description of datasets which were used to evaluate mm2-fast. Each of these were mapped to GRCh38 human genome reference.

| Query data set | Genome sample | Number of reads/contigs | N50 | Maximum length | Source |
| --- | --- | --- | --- | --- | --- |
| ONT | HG002 | 19M | 50K | 543K | https://precision.fda.gov/challenges/10/view |
|  | HG003 | 24M | 44K | 760K |  |
|  | HG004 | 29M | 48K | 1.1M |  |
| PacBio HiFi | HG002 | 8M | 13K | 30K | https://precision.fda.gov/challenges/10/view |
|  | HG003 | 7M | 15K | 32K |  |
|  | HG004 | 7M | 15K | 31K |  |
| PacBio CLR | HG002 | 30M | 11K | 89K | https://github.com/genome-in-a-bottle/giab_data_indexes |
|  | HG003 | 15M | 11K | 26M |  |
|  | HG004 | 13M | 10K | 5M |  |
| Genome assemblies | CHM13 | 24 | 154M | 248M | NCBI (GCA_009914755.3) |
|  | HG002 (hap1) | 523 | 46M | 107M | https://zenodo.org/record/4393631/files/NA24385.HiFi.hifiasm-0.12.hap1.fa.gz |
|  | HG002 (hap2) | 507 | 40M | 131M | https://zenodo.org/record/4393631/files/NA24385.HiFi.hifiasm-0.12.hap2.fa.gz |

### Minimap2 profile

We profiled a single-threaded execution of minimap2 using datasets listed in Table 1, and separately measured time consumed by three key modules (1) seeding, (2) chaining (DP chaining and RMQ-based DP chaining), and (3) alignment. Figure 2 shows the performance comparison and profile of minimap2 and our optimized implementation (mm2-fast). All runtime values shown are normalized by the total time consumed by minimap2 corresponding to each dataset. For profiling using a single thread, we used a random subset of 100K reads from each of the ONT, PacBio CLR, and PacBio HiFi datasets, but no sampling was done in the case of draft genome assemblies. We observed that the three modules collectively contribute to around 85% to 97% of the total mapping time across different datasets. Breakdown of time consumption among the modules was: seeding (3-13% time), chaining (9-68%) and alignment (18-76% time). Out of the time spent in chaining, 0-54% was spent in DP chaining, while RMQ based chaining accounted for 4-36% time. Interestingly, the time distribution of the three modules varied across all the input data types. For instance, the chaining was the most time consuming step for ONT and assembly datasets, whereas PacBio CLR and HiFi datasets required spending majority of the time in the alignment phase. Therefore, we focused on all the three key modules to achieve better performance.

**Figure 2.**
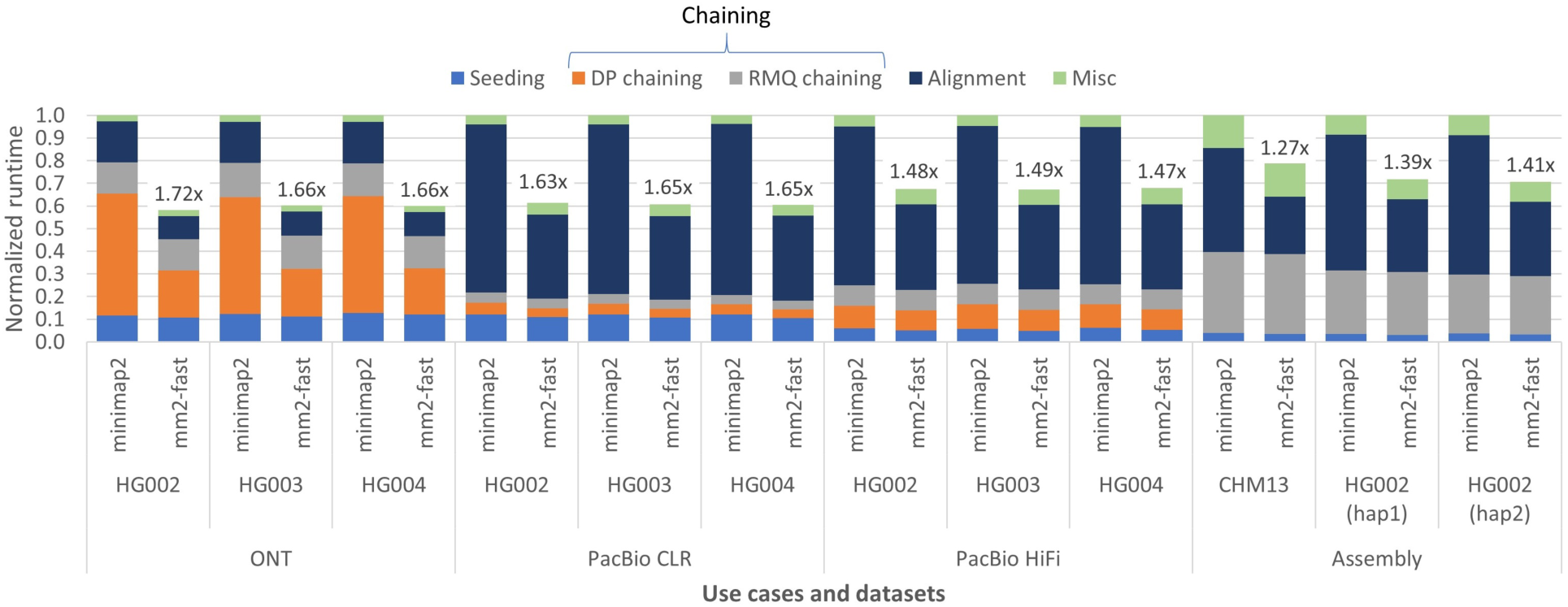
Work distribution for three modules: (1) hash look-up, (2) chaining (DP chaining and RMQ based chaining), and (3) alignment for minimap2 and mm2-fast across different datasets. Both the implementations were run using a single thread of Cascade Lake CPU. X-axis shows various query datasets, y-axis is the normalized time with respect to the mapping time consumed by minimap2 corresponding to each dataset. Speedup achieved by our mm2-fast over minimap2 for randomly sampled 100K reads for ONT, PacBio CLR, and PacBio HiFi and assembly contigs is shown on top of the mm2-fast bars.

### Summary of our optimizations

In mm2-fast, we implemented the following optimizations while ensuring that the mapping output obtained from our optimized minimap2 remains identical to minimap2:

- **Seeding**. We replaced hash-based minimizer lookup with learned index-based search over sorted list of the minimizers in the reference sequence. Internally, learned indexes use machine learning models to predict the positions of the desired minimizers. This resulted in nearly 3 − 4× speedup in minimizer lookup and up to 1.15× speedup in the seeding phase.
- **Chaining**. We accelerated DP chaining by vectorizing the traversal over the predecessor anchors using SIMD instructions and 32-bit integer/floating-point representation. Our AVX512-based vectorized chaining achieved up to 3.1 × speed-up over the implementation of DP chaining in minimap2.
- **Alignment**. Minimap2 implements base-level alignments using SSE2 instructions with 128-bit vector registers. As AVX2 and AVX512 instructions with support of 256-bit and 512-bit vector registers, respectively, are available in majority of modern general-purpose processors, we modified the alignment phase to add AVX2 and AVX512 based implementations. Our AVX512-based version yielded up to 2.2 × speed-up over the SSE2-based implementation in minimap2.

### Performance comparison

In Figure 2, the bars for minimap2 and our optimized implementation (mm2-fast) show relative time consumption of each module across various datasets using a single thread on Cascade Lake CPU. The speedups achieved for each dataset is also shown on top of the bars of mm2-fast. For single threaded execution, we achieved up to 1.7× speed-up compared to minimap2.

Figure 3 shows performance comparison of minimap2 and mm2-fast over full datasets listed in Table 1 using multi-threaded execution on an entire socket of Cascade Lake CPU. The labels above the bars for minimap2 show the end-to-end mapping time in hours, and the labels above the bars of our optimized implementation mm2-fast show the speedup achieved. Using multi-threaded execution on a single socket, we achieved up to 1.8 × speed-up compared to minimap2. mm2-fast also scales well with multi-threading. On a single socket system with 28 cores, we achieved up to 24.5× speed-up compared to single-threaded execution (Supplementary Figure S1). mm2-fast consumes nearly the same amount of memory as minimap2 (Supplementary Table S2). Step by step guide to use mm2-fast and verify the correctness is provided in Supplementary Note 1. The optimization details and the design choices are described in Methods section.

**Figure 3.**
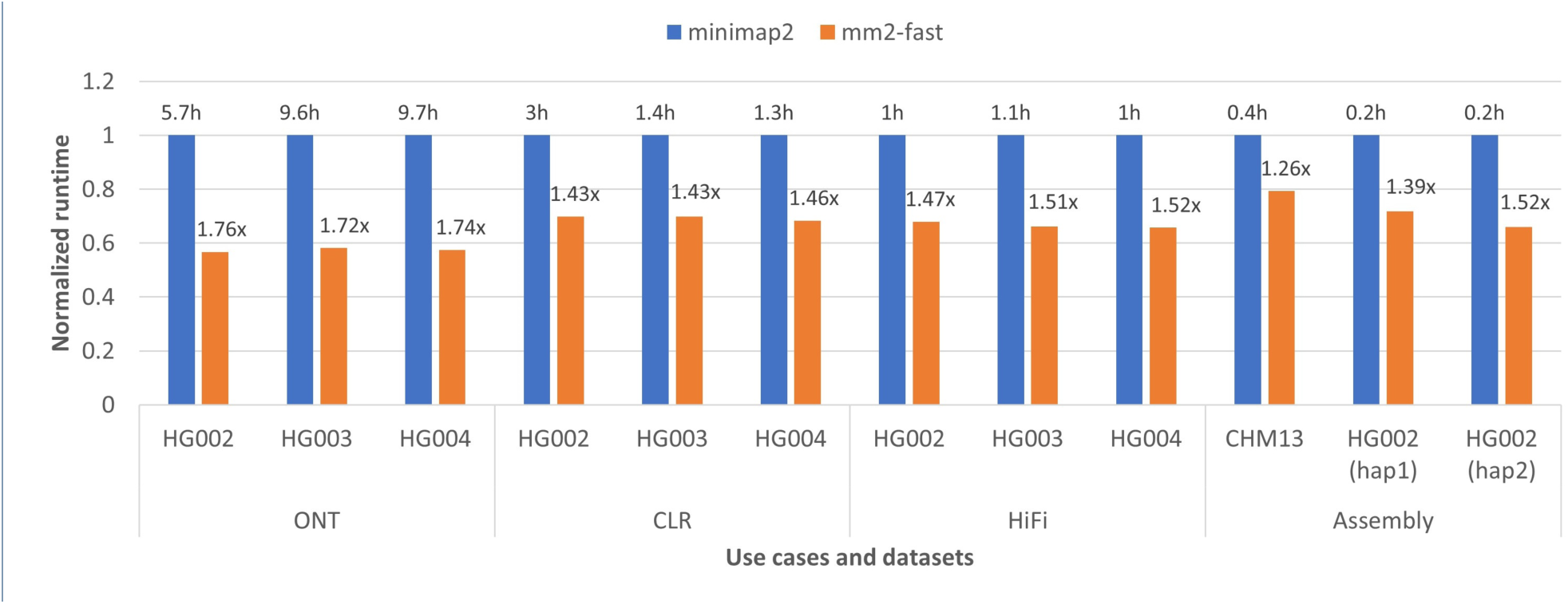
Performance comparison of minimap2 and minimap2-fast on a single socket Cascade Lake CPU (28 cores,) for full datasets. X-axis shows various query datasets, y-axis is the normalized time with respect to the mapping time taken by minimap2 corresponding to each dataset. On top of the bars of minimap2, we show the actual mapping time in hours by minimap2. The speedup achieved by minimap2-fast is shown on top of the bars of mm2-fast.

### Cross-platform performance and compatibility

To ensure that our optimizations deliver speedup across various architectures, we compared performance of mm2-fast and minimap2 on three generations of Intel architectures– (i) Skylake, (ii) Cascade Lake, and (iii) Ice Lake – and the recent AMD Rome architecture. The first three support both AVX2 and AVX512 vector processing, so we used AVX512 version of mm2-fast on them for these experiments. Rome only supports AVX2 and hence, AVX2 version of mm2-fast was used on Rome. Supplementary Table S1 provides the details of the architectural specifications of these systems. Figure 4 shows the speedups achieved on the four architectures. For each of the query datasets, we consistently achieved high speedups on Skylake, Cascade Lake, Ice Lake, and Rome processors. Note that these systems with different architectures run on different turbo-frequencies, so, their relative performance is not comparable.

**Figure 4.**
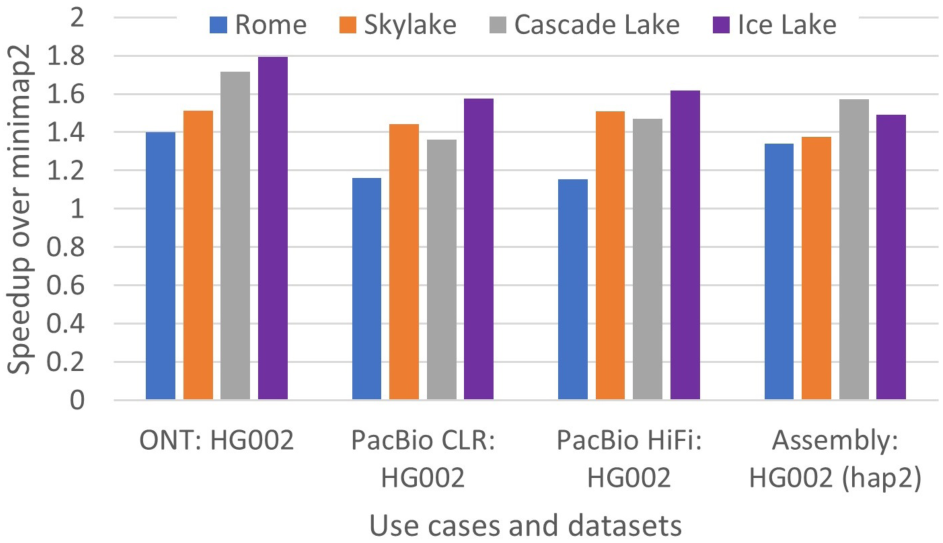
Cross-platform performance of our optimizations for Rome, Skylake, Cascade Lake and Ice Lake architectures using single socket. X-axis shows various query datasets and y-axis indicates the speedup achieved by mm2-fast over minimap2 – both running on the same CPU. The maximum speedup achieved was 1.8×.

### Time required for construction of the learned index

Across the four use-cases mentioned in Table 1, the construction of the learned index for mm2-fast takes only 2min23sec to 3min27sec. Moreover, the construction of the learned index is a one time activity for any reference sequence and use-case combination. Thus, the time spent in the construction of index gets amortized over the multiple samples that are mapped against the index. Therefore, we did not include it for both mm2-fast and minimap2 during the comparisons.

## Discussion

Improving long-read and genome assembly mapping time is important for three reasons: (a) it cuts down waiting time for a general user, (b) it is desirable for population-scale sequencing to achieve better throughput and reduce cloud computing costs, and (c) it improves the efficiency of real-time sequencing applications, including targeted sequencing [18–20], by matching speed of computation with the rate of datageneration.

Mapping tools designed for long-read analysis need to account for high sequencing error-rate, and as a result, involve complex heuristics to maintain scalability using large genomes. Three computational modules, i.e., seeding, co-linear chaining and alignment were identified to be the most time consuming steps in minimap2. We focused on accelerating all three of them while developing mm2-fast.

During the development of mm2-fast, we did extensive profiling of software performance, e.g., to optimize the count of instructions executed, cacheefficiency, and other important hardware parameters which dictate the CPU performance. *mm2-fast* is designed to achieve end-to-end hardware-aware acceleration of minimap2 on CPUs, while maintaining identical output. Unlike minimap2, mm2-fast implements a learned-index to allow faster minimizer look-ups, a SIMD-parallel co-linear chaining algorithm, and a revised implementation for sequence-alignment. Although the proposed optimizations will generally be useful for any tool which follows seed-chain-align procedure [21–23], we chose minimap2 to demonstrate the impact of these optimizations because it is a commonly used read mapper. mm2-fast leverages features available in modern CPUs (e.g., wide SIMD instructions), and will work on any modern general-purpose processor. mm2-fast offers a similar user-interface as minimap2 for compilation, learned-index-construction and read mapping.

Using mm2-fast, we achieved variable speedups depending on the type of input data, i.e., ONT, PacBio-CLR, PacBio-HiFi, and genome assemblies. This is attributed to the fact that input sequence lengths and error-rates change with the type of data, and also minimap2 uses different parameters (e.g., k parameter to set k-mer size) for each type of data. As output of mm2-fast remains identical to minimap2 (v2.22) in all scenarios, mm2-fast can be directly used as a faster alternative to minimap2. New features may be added in future into minimap2’s algorithm, at which point the output of mm2-fast and minimap2 may differ. To manage this better, we are working with the author of minimap2 to merge our optimizations in next release of minimap2. Currently, mm2-fast is made available as open-source code on GitHub as a branch in minimap2 repository https://github.com/lh3/minimap2/tree/fast-contrib-v2.22.

## Methods

### Seeding using a learned index

A recent work on learned index structures by Kraska *et al*. [24] has shown that an index structure can be viewed as a model that maps a key to its position, and therefore, can be replaced by machine learning models. For instance, a B-tree can be seen as a model that maps a key to its position in a sorted list. The learned index structures can take advantage of the distribution of the keys and train a machine learning model, e.g., a recursive model index (RMI), such that it outperforms traditional B-trees in search time and memory footprint. Following [24], several other learned index structures have been proposed [25–28]. Learned indexes have emerged as a performant alternative to solve problems in various domains including bioinformatics, e.g., for genome indexing using FM-index [14] and suffix array [29]. In this work, we design a learned index based hash table to improve the query time for minimizer lookup during the seeding phase.

### Hash table implementation in minimap2

A hash table index is used in minimap2 to store all minimizers of a reference sequence as keys and their positions in the reference sequence as values. In the seeding phase, a successful hash lookup returns all positions where a query minimizer occurs in the reference sequence. Typically, such hash table lookups are known to be faster. However, they may incur overheads due to a large number of collisions, and also incur high cache misses as a side-effect of irregular memory accesses during hash table access, including, collision resolution. For instance, the hash table in minimap2 uses a traditional quadratic probing technique which may lead to sub-optimal cache performance. In the following, we present our learned-index-based hash table design which outperforms the traditional hash table implementation in minimap2.

### Design of a learned hash table

Hash lookup problem can be modeled as a search on a list of key-value pairs that is sorted according to the keys. In our learned hash table design, we maintain two data structures as illustrated in Figure 5. The first is a key-sorted list of key-value pairs where keys are the actual minimizers. The second is a `position-list` containing a concatenated list of lists of positions of all the minimizers. For each minimizer key, the value part of the first data structure encodes the starting index in the `position-list` and the count of the positions for the minimizer. For instance, in Figure 5, a minimizer entry: *mm*5 → [8, 3] represents that *mm*5 appears 3 times on the reference sequence – at index 5, 21, and 57; and these three positions are stored at three consecutive locations in `position-list` starting from index 8. We use a learned index structure to search for the minimizer entry in the first data structure.

**Figure 5.**
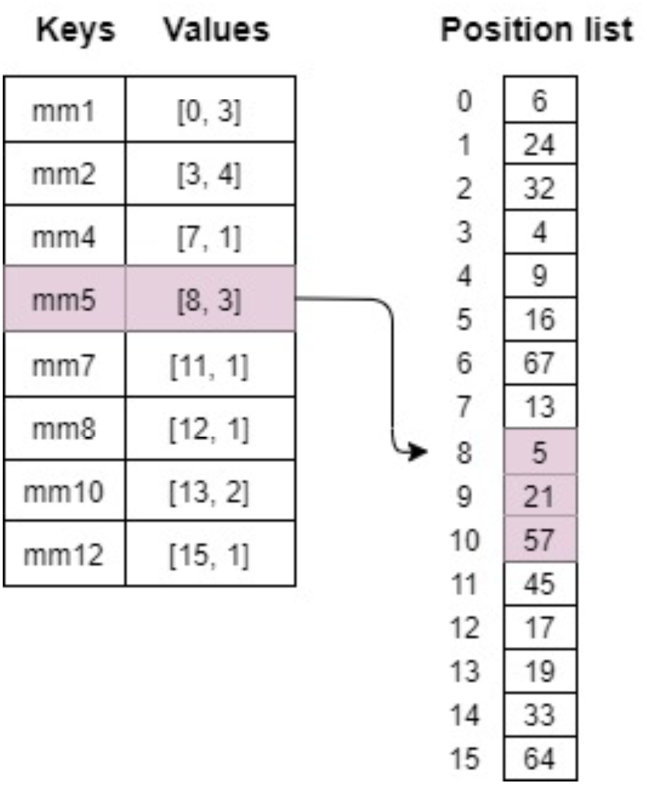
Data structures used for hash table: Minimizers extracted from the reference sequence are stored in a sorted list as key-value pairs. Position list maintains a separate list of the positions of minimizers on the reference sequence.

Among the proposed learned index structures, RMI exhibits the best performance/size tradeoff for real-world read-only in-memory dense arrays [30]. RMI consists of a multi-layer tree structure with a model at each tree node. An RMI with more than two layers is almost never needed [31]. Therefore, we train a two-layer RMI model to learn the distribution of the sorted keys and use the trained RMI to search through the sorted list. Figure 6 shows a two-level RMI model and illustrates, with an example, how we perform a lookup operation. While performing a lookup operation for a minimizer, the model at the root layer is used to predict the correct model to use at the leaf layer. The predicted model at the leaf layer is used to predict the position of the key in the sorted list. RMI guarantees that the key is present within a certain range from the predicted location [30]. If the desired key is not found at the predicted position, the last mile search is conducted in the provided range to find the key. The last mile search is typically short because the key is expected to be in proximity of the predicted location. In our experiments, we observed that RMI-based lookup is nearly 3 − 4 × faster than the existing hash table implementation in minimap2.

**Figure 6.**
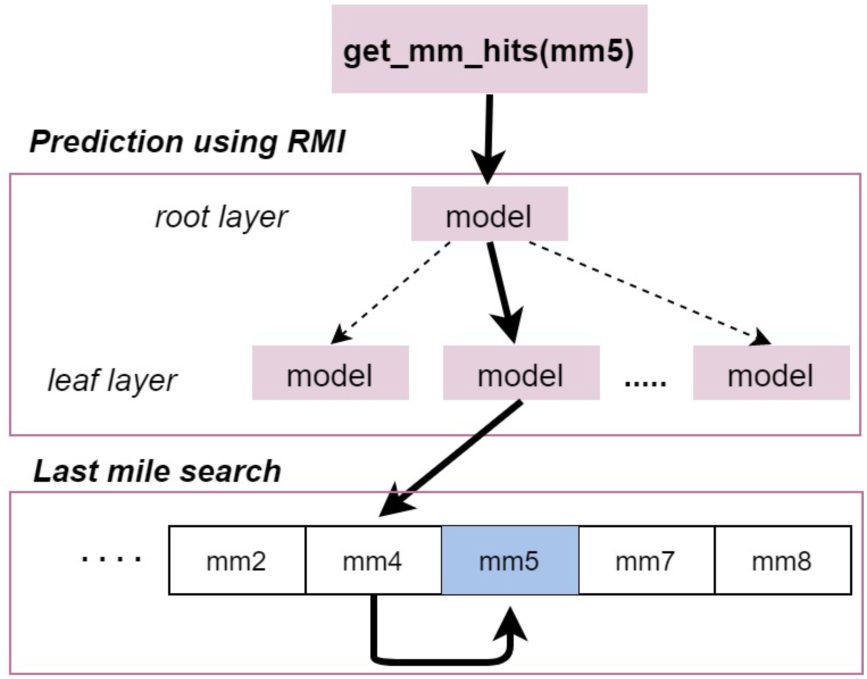
Two-layer RMI: An example minimizer lookup is illustrated - *get_mm_hits*(*mm*5) calls a lookup for a minimizer *mm*5. The RMI root predicts the leaf layer model which in turn predicts the location of *mm*4 in the sorted list. Finally, the last mile search from *mm*4 walks to the location of *mm*5 and returns its value to the caller.

### Design choices

We make the following design choices and apply architecture-aware optimizations similar to [14] so as to achieve maximum speedup using RMI:

- **Number of leaf nodes**. The number of leaf layer models plays a crucial role in the efficiency of the RMI lookup [30]. Using a large number of leaf layer models delivers better prediction and shorter last-mile searches - at the cost of larger memory consumption. By default, we use *n/*32 leaf layer models where *n* is the total number of minimizers in the list since empirically that provided a good trade-off between prediction accuracy and memory consumption.
- **Vectorized last mile search**. The last mile search is performed using binary search. Once only a few elements remain for search (say, less than eight 64-bit elements for AVX512 vector instructions), we compare them simultaneously using SIMD instructions.
- **Batched processing**. Irregular memory accesses while traversing through the RMI tree and the last mile search lead to higher cache misses, which adversely affect the performance. We use software prefetches to hide the memory latency by processing a batch of lookups at a time. Each individual lookup can be split into a sequence of steps to be performed - (i) visit the RMI root and predict leaf layer model, (ii) visit leaf model and predict the location of the key, and (iii) one step for each iteration of the binary search performed around the prediction. During batch processing, we process all the lookups in a batch in round-robin fashion. Every time a lookup gets a turn, we advance that lookup by one step and use software prefetches to start prefetching the data into the caches for its next step. While the data is getting prefetched, we go over the rest of the lookups in the batch one by one and advance them by one step and start their prefetches. By the time, a lookup gets a turn again, the data it requires for the next step is already expected in cache. If all the steps of a lookup are done, we replace it with the next unprocessed lookup from outside the batch. We continue this till all the lookups are done. The batch size should be large enough to hide the memory latency but not so large that data corresponding to all the lookups in a batch do not fit in cache. We theoretically compute a range of ideal batch sizes and empirically find the best batch size from that range.

### SIMD acceleration for co-linear chaining

#### Chaining in minimap2

The seeding phase outputs a list of all identified anchors sorted according to their position in the reference sequence for further processing in the chaining stage. An anchor is defined as a matching minimizer between a query and a reference sequence. It can be represented as a 3-tuple (*r, q, l*), where *r* and *q* are the positions of the matching minimizer on the reference and the query sequence, respectively, and *l* is the length of the minimizer.

Given a list *L* :{*a*_1_, *a*_2_,…, *a*_*n*_} of anchors sorted according to their reference positions, the chaining step identifies ordered subsets of co-linear anchors as chains which achieve the highest chaining scores. Let *S*(*i*) be the highest chaining score for a chain that ends at anchor *a*_*i*_; *S*(*i*) is computed using the following dynamic programming (DP) recursion in minimap2:

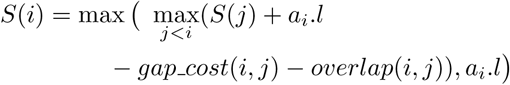

where *gap cost*(*i, j*) function penalizes the score based on the distance between anchors *a*_*i*_ and *a*_*j*_, and *overlap*(*i, j*) denotes the count of overlapping bases between anchors *a*_*i*_ and *a*_*j*_. DP chaining linearly scans the previous *S*(*j*) values, thus requiring the *O*(*n*) time in computation of every *S*(*i*). RMQ based DP chaining performs an RMQ query over a binary tree to get the best *S*(*j*) in *O*(*log*(*n*)) time, but needs to simplify the cost function to enable that. In mm2-fast, we accelerate the former. Figure 7 depicts the chaining of two co-linear anchors and their corresponding gaps and overlap.

**Figure 7.**
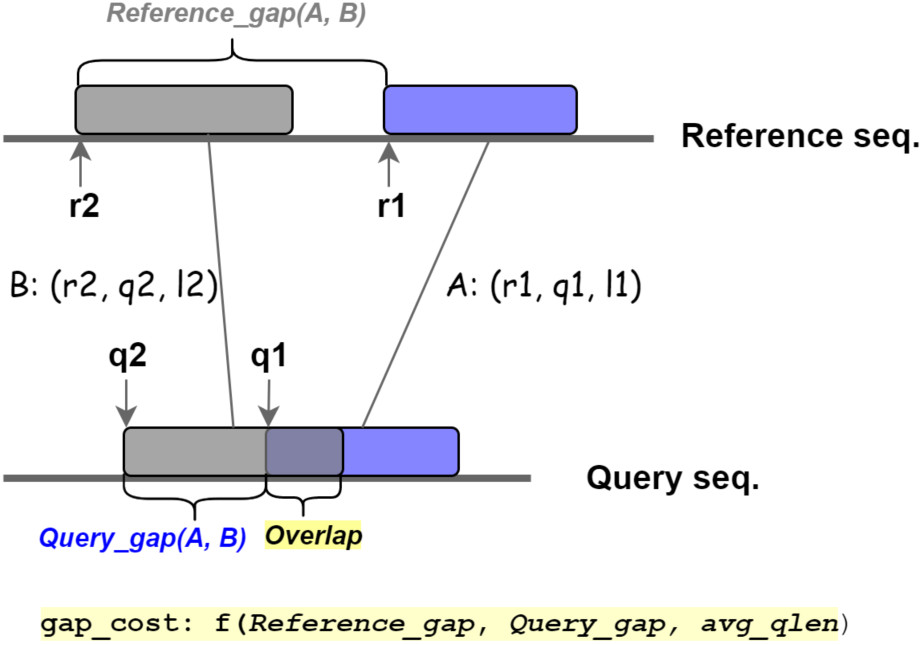
Chaining of two co-linear anchors *A* and *B*. Here two anchors overlap on the query sequence. Gap cost function in minimap2 is calculated using the reference gap, query gap, and the average length of all anchors *avg_qlen*

#### Dynamic Programming (DP) chaining in minimap2

Algorithm 1 presents a DP-based anchor-chaining implementation in minimap2. For each anchor *a*_*i*_ in the list (line 3), the inner loop (line 8) iterates over all the anchors *a*_*j*_ where *start ≤ j < i*. DP-based execution pattern ensures that the maximal chaining scores till anchor *a*_*i-*1_ are already computed before the computation of score for *a*_*i*_. The expression in line 11 computes the chaining score. The two variables *max score* and *predecessor index* (lines 12-14) track the maximum scoring chain found so far and the index of the predecessor anchor connected to it. Considering that the gap cost and the overlap can be computed in constant time, the worst-case time complexity for the whole chaining step is *O*(*n*^2^).

Minimap2 applies a set of heuristics to accelerate the chaining step. The `while` loop (line 6) decides the range of predecessors based on the distance between the two anchors with respect to the reference sequence (i.e. Reference gap, as illustrated in Figure 7). For instance, if the gap between two anchors is above a user-specified threshold, the gap cost is assumed to be ∞. As the anchors are sorted, the range of anchors with larger distance can trivially be ignored for the score computation. The `if` condition in the inner loop (line 9) applies another set of filters based on the distances between the anchors with respect to the query sequence (i.e. Query gap, as illustrated in Figure 7). The anchor being considered is ignored if the gap is above a certain threshold or negative. After every iteration of the inner loop, the *max skip* condition (line 15) is evaluated. According to the heuristic, if we do not find a better score over the last *max skip* attempts, then the inner loop terminates. In minimap2, the default value for *max skip* is 25. The sole purpose of this heuristic is to accelerate the chaining step potentially at the cost of chaining accuracy. The heuristic can be disabled by setting *max skip* to a large value (say ∞) to achieve better chaining accuracy. The actual conditional expressions and further implementation details can be found in minimap2 paper [7].

##### Algorithm 1 Sequential chaining as used in minimap2

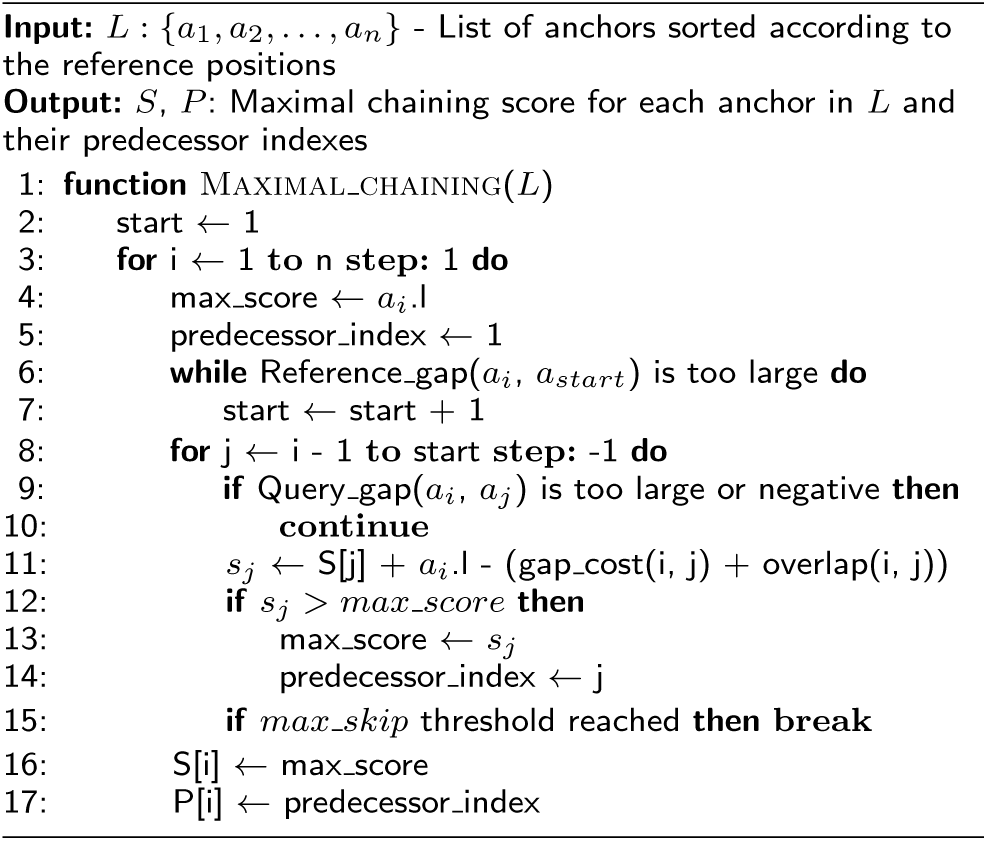

#### Our SIMD based DP chaining

As shown previously in Results section, chaining is one of the most compute-intensive steps in the complete pipeline. Despite various heuristics used in the chaining step, the majority of the time is consumed in the execution of the inner loop. We accelerate the chaining step by redesigning the inner loop to exploit SIMD parallelism. Typically, modern compilers provide support for auto-vectorization using compilation flags or annotations which can identify opportunities to convert sequential loops to their SIMD version. However, in practice, a loop with dependencies and branching instructions makes auto-vectorization difficult. We tried auto-vectorizing the inner loop using the latest versions of gcc and icpc compilers and verified that none of the modern compilers could auto-vectorize the loop. Designing a SIMD-friendly chaining algorithm is nontrivial because the inner loop contains multiple conditional branches, such as *Query gap* and *max skip*, and during the computation of *max score*. In our optimizations, we carefully resolved the branches and inter-loop dependencies and adopted a hybrid (sequential+SIMD) approach to maximize the vectorization gains. We ensured that the sequential and vectorized versions produce identical chaining output.

##### Algorithm 2 Proposed vectorized chaining

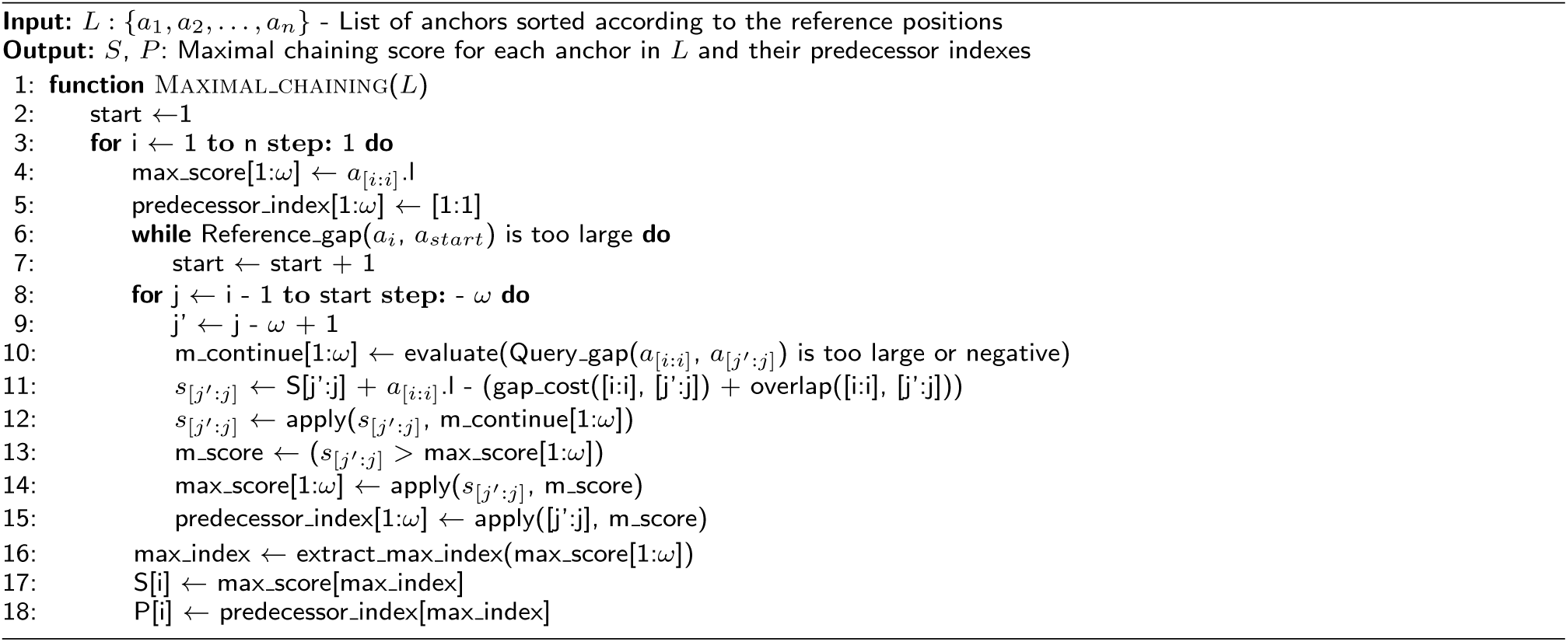

More specifically, we made the following improvements to the chaining module

- **Inner loop vectorization**. Algorithm 2 shows our vectorized algorithm for anchor chaining using 32-bit number representation. We denote *ω* as a SIMD-width. For instance, using AVX512 bit instructions and 32-bit numbers, we can process 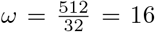 elements simultaneously. The inner loop at line 8 implements a *ω*-way vectorized version that computes the maximal chaining scores. Vector instructions perform the same operation over multiple data elements. In Algorithm 2, the notation A[i:j] represents a vector operation on a range of data elements A[i] to A[j]. A[i:i] broadcasts the value of A[i] to the vector register. Similarly, [i:j] and [i:i] loads and broadcasts the constant values to the registers respectively. Conditional branches (`if` conditions) are vectorized using *ω*-bit vector masks; if the branch is taken, the respective bit in the vector mask is set to 1 (line 10). We vectorized *gap cost* and *overlap* evaluation, and compute the chaining scores for *ω* predecessor anchors in one vectorized iteration. Once the chaining score is computed, we apply the computed mask *m continue* (line 12) such that the scores are zeroed out when the respective bit is set in the mask. Similar to the scalar version, the two vector registers, *max_score* and *predecessor_index*, track the maximum scores across vector lanes. After every loop iteration, the *max_score* is compared against the computed chaining scores and masked so that the maximum score is guaranteed to be present in the register. Finally, we extract the maximum score and the predecessor index using a sequential pass over *max_score*. (line 16).
- **Hybrid vectorized+sequential execution**. In an ideal scenario, loop vectorization delivers maximum benefit when there is no path divergence due to conditional instructions. In the presence of path divergences, vectorized implementation has to compute all control paths and use appropriate masks to obtain the desired outputs. For instance, we need to compute the chaining scores for all vector lanes even though some of the iterations would have continued without computing the chaining scores in the sequential algorithm (Algorithm 1, line 10). This results in wasted computation over the vector lanes and leads to sub-optimal performance. Using AVX512 vectorization, out of 16 vector lanes, we typically found sufficient inner loop iterations computing the score, thus, benefiting from the optimizations. However, in the case of shorter inner loops (say, *<* 16 iterations), we observed that only a few iterations in the sequential algorithm computed the chaining score. In such a case, due to path divergence, a single vector-iteration might incur more instructions than the sequential executions of *few* scalar iterations. We mitigated this issue by adopting a hybrid approach: mm2-fast falls back to the sequential execution of the inner loop if there are fewer than five iterations; else, it uses the vectorized implementation.
- **Disabling max skip heuristic**. As mentioned earlier, *max_skip* condition in Algorithm 1 uses a heuristic parameter *max_skip*, which accelerates chaining in minimap2 at the cost of chaining accuracy. Moreover, *max_skip* condition also poses challenges to vectorization as it carries loop dependencies. In our optimizations, we disabled this heuristic by setting *max_skip* to ∞ and removed the condition from our vectorized implementation. This benefited us in two ways: first, we resolved the loop carrying dependency resulting in less complex SIMD implementation and better speedup, and second, we achieved better chaining accuracy. Note that the performance improvement reported in the result section compares the performance of our optimizations with *max_skip* = ∞ against minimap2 with its default setting *max_skip* = 25. We achieve up to 3.1× speedup in chaining over minimap2 with default heuristics *max_skip* = 25, and up to 8.4× as compared to minimap2 with *max_skip* = ∞. Supplementary Figure S2 shows the end-to-end performance comparison of mm2-fast against minimap2 with *max_skip* = 25 and *max_skip* = ∞.

### SIMD acceleration for pairwise sequence alignment

Minimap2 computes DP-based global sequence alignment to extend through the gaps between adjacent chained anchors. The presence of long gaps between anchors results in slower DP-based alignment, which can be accelerated using SIMD-based vectorization. However, longer sequences for alignment demand more bits to capture the alignment score; this leads to a lower number of available vector lanes and hence lower parallelism. Minimap2 adopts the Suzuki-Kasahara (SK) formulation for DP-based alignment. SK formulation bounds the number of bits required to capture the score in a DP matrix to the *scoring parameters*. In practice, these bounds ensure that 8-bits are sufficient for maintaining score values. Therefore, with 128-bit vector processing available with SSE, 16-way parallelism is available.

Minimap2 applies intra-task parallelism, exploiting parallelism in a single DP matrix. All the DP matrix cells along an anti-diagonal are independent of each other. Thus, minimap2 applies vectorization with SSE SIMD instructions along anti-diagonals to accelerate the alignment. To do so, minimap2 switches from row-column-based matrix coordinates to diagonal-anti-diagonal-based coordinates. Additionally, to reduce the computation burden, minimap2 restricts the DP cell computations to a certain band around the main anti-diagonal. Minimap2 uses a default band value of 500. Precise details about the alignment computation are available in [7, 32].

SSE instructions provide only 16-way parallelism. Using 512-bit vector processing (AVX512), the available parallellism increases 4-fold to 64-way. Moreover, the default band value in minimap2 is large enough to keep all the vector lanes of AVX-512 occupied. In mm2-fast, we accelerated the DP-based alignment by utilizing AVX-512 vectorization and used additional logic to maintain the output identical to minimap2. To support a wide range of processors, we also developed an AVX2 version.

Manymap software [10] also provides an accelerated version of the alignment module of minimap2 by upgrading the vectorization to AVX-512. Manymap also applies optimizations to marginally reduce the instructions in the inner DP loop. However, its alignment output differs from minimap2. In minimap2, the actual band of DP matrix that is computed could be greater than or equal in size to the given band size; this is because actual anti-diagonal computed in minimap2 is a multiple of SIMD width (8 for SSE). SIMD widths change when we switch from SSE to AVX2/AVX512. Consequently, the same DP matrix computations with longer SIMD widths can result in computing a bigger band size. Therefore, to maintain the exact same output, we introduced additional instructions in our implementations to ensure the computation of the exact same band as minimap2. Manymap does not guarantee the same (Supplementary Note 2). Due to this, we skip comparing the performance of mm2-fast with Manymap.

Ideally, moving from SSE to AVX-512 has a potential of 4× improvement in the runtime. However, in practice, we saw 1.8 − 2.2× speedups due to the following factors: (a) smaller sequences lead to underutilization of vector lanes, (b) smaller anti-diagonals near the two corners of DP-matrix lead to idle vector lanes, (c) latency and throughput difference between SSE and AVX-512 vector instructions, and (d) use of additional instructions in the AVX-512 version to ensure the exact same output as the SSE version. The memory requirements and access pattern of the AVX2 and AVX512 versions in mm2-fast remain the same as the SSE version in minimap2.

## Availability of data and materials

mm2-fast source code is available under the open source MIT license at https://github.com/lh3/minimap2/tree/fast-contrib. Datasets used for benchmarking are publicly available (Table 1).

## Ethics approval and consent to participate

Not applicable.

## Authors’ contributions

SK led the software implementation of mm2-fast. All authors contributed to design of algorithm, experiments and manuscript preparation. All authors read and approved the final manuscript.

## Optimization Notice

Software and workloads used in performance tests may have been optimized for performance only on Intel microprocessors. Performance tests, such as SYSmark and MobileMark, are measured using specific computer systems, components, software, operations and functions. Any change to any of those factors may cause the results to vary. You should consult other information and performance tests to assist you in fully evaluating your contemplated purchases, including the performance of that product when combined with other products. For more information go to http://www.intel.com/performance. Intel, Xeon, and Intel Xeon Phi are trademarks of Intel Corporation in the U.S. and/or other countries.

