## Supplementary Material for "Accelerating long-read analysis on modern CPUs"

### Supplementary Data

**Table S1** Architectural specifications of the four processors: Skylake, Cascade Lake, Ice Lake and Rome which were used for the experiments.

| | Intel® Xeon® Platinum 8180 Skylake | Intel® Xeon® Platinum 8280 Cascade Lake | Intel® Xeon® Platinum 8380 Ice Lake | AMD EPYC™ 7742 Rome |
|---|---|---|---|---|
| Sockets × Cores × Threads | $1 \times 28 \times 2$ | $1 \times 28 \times 2$ | $1\times$ 40 $\times 2$ | $1 \times 64 \times 2$ |
| AVX register width (bits) | 512, 256, 128 | 512, 256, 128 | 512, 256, 128 | 256, 128 |
| Vector Processing Units (VPU) | 2/Core | 2/Core | 2/Core | 2/Core |
| Base Clock Frequency (GHz) | 2.5 | 2.7 | 2.4 | 2.25 |
| L1D/L2 Cache (KB) | 32/1024 | 32/1024 | 48/1280 | 32/512 |
| L3 Cache (MB) / Socket | 38.5 | 38.5 | 60 | 256 |
| DRAM (GB) / Socket | 96 | 96 | 128 | 128 |
| Bandwidth (GB/s) / Socket | 112 | 128 | 205 | 205 |
| Compiler Version | ICPC v. 19.1.3.304 | | | |

**Table S2** Memory-consumption (GB) of minimap2 and mm2-fast evaluated with various datasets using 28-core multi-threaded execution.

| Query dataset | minimap2 | mm2-fast |
|---|---|---|
| ONT: HG002 | 33.5 | 34.1 |
| PacBio CLR: HG002 | 23.2 | 23.5 |
| PacBio HiFi: HG002 | 28.1 | 30.3 |
| Assembly: HG002 (hap2) | 28.9 | 29.3 |

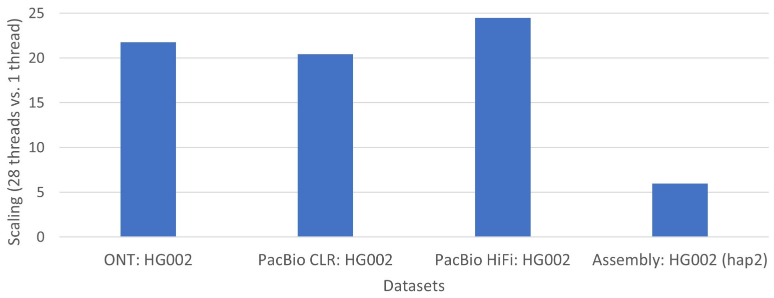


**Figure S1** Speedups achieved using multi-threaded execution of mm2-fast using single socket with 28 cores. X-axis shows query datasets used, and y-axis shows the ratio of single threaded execution time and 28-core execution time.

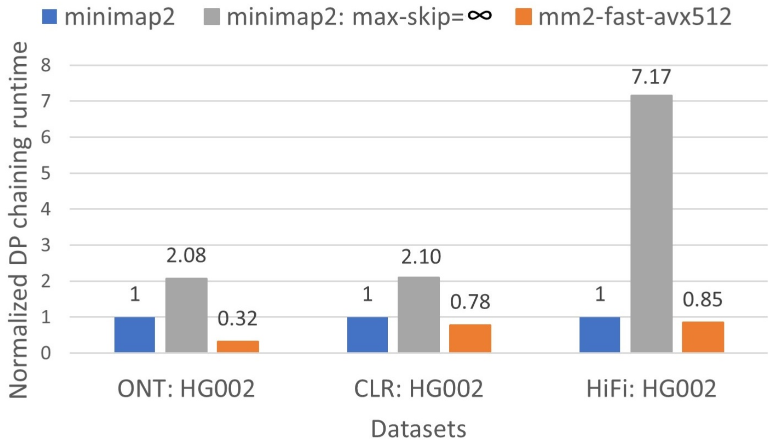


**Figure S2** Performance comparison of dp chaining of mm2-fast and minimap2 as well as a version of minimap2 in which its $max\_skip$ chaining heuristic is disabled by setting $max\_skip$ parameter to $\infty$. X-axis shows various datasets used, and y-axis is the normalized time with respect to the dp chaining time taken by minimap2. Minimap2 slows down by up to $7.17\times$ (HiFi dataset) when its $max\_skip$ heuristic is turned off. Assembly use case does not use DP chaining and hence, is excluded from this experiment.

#### Supplementary Note1: Correctness check

We ensure that minimap2 (v2.22) and mm2-fast produce identical output. Minimap2 uses *max_skip* heuristic to speed up the performance at the cost of the chaining accuracy. The default minimap2 configuration uses *max_skip*=25. For better accuracy, *max_skip* can be set to a higher value using the command-line flag `--max-chain-skip`. A larger value of *max_skip* heuristic provides better mapping accuracy. We do not use the *max_skip* heuristic in our vectorized DP chaining implementation. Therefore, the output of mm2-fast should match the most accurate mapping output of minimap2 for DP chaining, i.e., with *max_skip* heuristic disabled. For verifying the correctness of mm2-fast, minimap2 should run with `--max-chain-skip=`$\infty$. For applying SIMD, mm2-fast needs to use one of the following compiler flags – `-march=native or -mavx2`. These flags result in additional automatic compiler optimizations – in parts of the source code that we have not manually improved – to produce more efficient assembly code. This results in the slightly different output - output CIGAR is the same, but some of the scores are off by 1. The same compiler flags added to the minimap2 compilation result in identical changes to the output.

Following are the steps to verify the correctness of mm2-fast. The example commands below use sample filenames *ref-seq* and *read-seq* for a reference sequence and a read sequence files respectively, and *map-ont* as a preset parameter.

**Clone minimap2 (v2.22)**:

```
git clone https://github.com/lh3/minimap2.git -b v2.22
```

**Compile and run**:

```
cd minimap2
```

Compile and run on the same machine. Add the following line to the Makefile after line 2:

```
CPPFLAGS+= -march=native
make
./minimap2 -ax map-ont ref-seq read-seq --max-chain-skip=1000000 > minimap2_output
```

**Clone mm2-fast**:

```
git clone --recursive https://github.com/lh3/minimap2.git -b fast-contrib-v2.22 mm2-fast-contrib
```

**Compile**:

Compile and run on the same machine.

```
cd mm2-fast-contrib && make multi
```

The above command should generate three executable files: 1. `mm2-fast` 2. `mm2-fast-lhash` 3. `mm2-fast-no-opt`. By default, `mm2-fast` applies two optimizations, AVX512 based chaining and AVX2/AVX512 based alignment. On top of these two optimizations, `mm2-fast-lhash` uses learned hash tables. The optimizations in mm2-fast require architectural support of AVX2/AVX512. In the absence of AVX2/AVX512, `mm2-fast-no-opt` can be used to run with all optimizations turned off.

##### Correctness check with `mm2-fast`

**Run** `mm2-fast`

```
./mm2-fast -ax map-ont ref-seq read-seq --max-chain-skip=1000000 > mm2-fast_output
```

**Match output files**

```
diff minimap2_output mm2-fast_output > diff_result
```

The file `diff_result` should show a clean-diff with the difference of 2 lines, i.e., the lines containing the command-line parameters for minimap2 and mm2-fast.

##### Enabling learned hash tables

To make the correctness verification seamless, by default, we have disabled learned hash tables as it requires an additional installation. Learned hash-table uses an external training library that runs on *Rust*. Following are the steps to enable learned hash table optimization in mm2-fast:

- Install Rust and add installation path to *.bashrc* file. This is fairly quick and can be done by a single command given at https://rustup.rs/.
- Create learned hash table index for a reference sequence and a preset parameter (say map-ont).
  `./build_rmi.sh ref-seq map-ont`
  Index building is one-time task for a reference sequence and a preset parameter, and can be reused for all subsequent executions. Note that, for a given reference sequence, the hash index changes with difference preset parameters.

- Once the index is built, run `mm2-fast-lhash`.
  `./mm2-fast-lhash -ax map-ont ref-seq read-seq > mm2-fast-lhash_output`
  The output file `mm2-fast-lhash_output` should also be identical to `minimap2_output` file produced above.

Supplementary Note2: Correctness check of Manymap

mm2-fast fulfills the criteria of the same output as Minimap2; thus, we put any competitive software through the same criteria before conducting a performance comparison. Manymap accelerated the pairwise sequence alignment module of Minimap2; however, our experiments show that it does not fulfill the exact output criteria. Since Manymap is based on Minimap2 v2.16, we used this version to perform the output test on Manymap. We used randomly sampled 100K queries from PacBio HiFi data. We experimented as follows::

**Run** `Manymap`

```
git clone https://github.com/RapidsAtHKUST/manymap.git manymap
cd manymap && make avx512=1
./minimap2 -ax asm20 ref-seq read-seq > manymap_output
```

**Run** `Minimap2-v2.16`

```
git clone https://github.com/lh3/minimap2.git -b v2.16
cd minimap2 && make
./minimap2 -ax asm20 ref-seq read-seq > minimap2-v2.16_output
```

The diff of `manymap_output` and `minimap2-v2.16_output` resulted in more than 20 lines of difference.
